# Breaking through evolutionary constraint by environmental fluctuations

**DOI:** 10.1101/016790

**Authors:** Marjon GJ de Vos, Alexandre Dawid, Vanda Sunderlikova, Sander J Tans

## Abstract

Epistatic interactions can frustrate and shape evolutionary change ^1, 2^ ^3–7^. Indeed, phenotypes may fail to evolve because essential mutations can only be selected positively if fixed simultaneously ^5,8,9^. How environmental variability affects such constraints is poorly understood. Here we studied genetic constraints in fixed and fluctuating environments, using the *Escherichia coli lac* operon as a model system for genotype-environment interactions. The data indicated an apparent paradox: in different fixed environments, mutational trajectories became trapped at sub-optima where no further improvements were possible, while repeated switching between these same environments allowed unconstrained adaptation by continuous improvements. Pervasive cross-environmental trade-offs transformed peaks into valleys upon environmental change, thus enabling escape from entrapment. This study shows that environmental variability can lift genetic constraint, and that trade-offs not only impede but can also facilitate adaptive evolution.

---

It is widely believed that epistatic interactions can direct evolutionary change ^1^. Epistasis has been implicated in shaping RNA ^10^ and protein ^4, 6, 7^ sequences, sensing ^5^ and translation ^11^ functions, as well as developmental programs ^12^ and speciation ^13–15^. Indeed, phenotypes may fail to evolve not because they are impossible biochemically or physically, but because essential mutations are mutually dependent, and must be fixed simultaneously to be selected positively ^5, 8, 9^. How the constraining effects of such genetic interactions are affected by environmental variability remains poorly understood. It has been shown that mutational effects 16-19 and epistasis itself ^20, 21^ can depend on the environment, that bacterial resistance evolution can be contingent on the rate of antibiotic increase ^22^, and that adaptation *in silico* can be accelerated by environmental change ^23–25^. These observations suggest that environmental variability may not only produce variable selection, but could also control phenotype accessibility and stasis.

To investigate how environmental variability affects genetic constraints, we focused on a model system for genotype-environment interactions, the *lac* regulatory system of *E. coli*. Its physiology has been studied extensively: in the presence of lactose, expression of the *lac* genes allows *E. coli* cells to import and metabolize lactose, while in the absence of lactose, repression of these genes limits physiological costs ^26, 27^. The ability to regulate *lac* expression relies on the binding of the *lac* repressor to the *lac* operator DNA upstream of the coding region. We surmised that the co-evolution of such protein-DNA interfaces could be constrained by epistatic interactions. In lock-key recognition, mutating either lock or key is expected to lead to recognition loss ^2, 9^. At the same time, mutating both lock and key may produce a different, better-matching pair. Indeed, the *lac* transcription factor phylogeny suggests extensive historic adaptation of the repressor-operator interface, and reveals multiple homologous repressors that bind specifically to their cognate operator ^28, 29^.

Mutational analysis of the *lac* repressor-operator interface has shown that two repressor residues and four operator bases control binding specificity (Fig. 1a and b) ^30, 31^. We constructed repressor and operator variants in these sites, and assayed them in the two contrasting conditions of *lac* regulation. We quantified the ability to repress the *lac* genes (*R*) as the inverse of the measured *lac* expression level in the *absence* of inducing ligand (Fig. 1c, Methods). The ability to express the *lac* genes (*E*) was quantified by the measured *lac* expression level in the *presence* of ligand (Fig. 1c). Note that the repression ability (*R*) is thus not the inverse of the expression ability (*E*). We identified two repressor-operator pairs, denoted as MK:acca and YQ:tggt (Fig. 1c), for which the fold-changes between the induced and non-induced expression level *R*•*E* were substantial (6 and 55 respectively), with *E* being approximately equal but *R* about 20-fold lower for MK:acca. The MK:acca genotype is thus able to regulate *lac* expression, but can improve repression ability by mutating the repressor (MK to YQ) and operator (acca to tggt).

**Figure 1.**
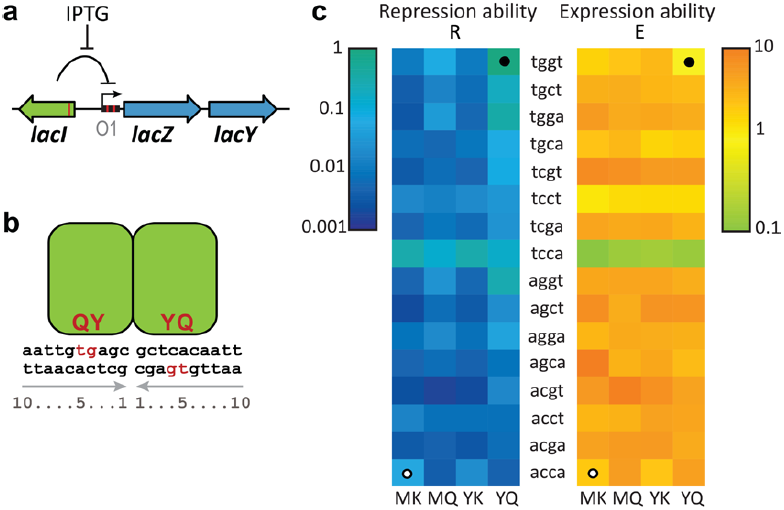
Repression and expression ability of *lac* repressor-operator mutants. (a) Schematic representation of the *Escherichia coli lac* system. β-galactosidase (LacZ) and the *lac* permease (LacY) are co-regulated by the repressor LacI. Expression is induced by isopropyl-β-d-1-thiogalactopyranoside (IPTG). Red lines correspond to mutated positions. (b) Mutated positions, responsible for specific repressor-operator binding in red. (c) Characterization of the 64 *lac* repressor-operator variants. The starting and final sequences are indicated by open and filled circle respectively. Repression ability (*R*) is the inverse of the measured expression level in the absence of IPTG. Expression ability (*E*) is the measured expression level in the presence of IPTG (Methods).

We investigated the interaction between non-cognate pairs by swapping around the two operators. *R* was found to be low for MK:tggt and YQ:acca (100 to 200-fold lower than for the cognate pair YQ:tggt). These data were consistent with the reciprocal sign epistasis hypothesised for lock-key interactions: exchanging either of the binding partners leads to binding loss, but changing both can restore it. This notion was supported by the overall expression levels for the intermediate MK:tggt and YQ:acca, which were high and unresponsive to ligand (*R*•*E* = 1). However, while the presence of reciprocal sign epistasis is required, it is not sufficient to constrain phenotypes on sub-optima ^9^. Indeed, the repressor and operator modifications both involve multiple mutations, and their one-by-one fixation in particular order ^1^ could confer continuous improvements in repression ability.

To test trajectories for all possible orders of all essential mutations, we constructed the remaining intermediate genotypes between MK:acca and YQ:tggt. In total, 720 trajectories can be taken along the 2^6^ = 64 genotypes. Analysis showed that all trajectories contained depressions in both *R* and *E* (Fig. 2, Supplementary Fig. 1). The depressions were at least 2 mutations wide and peaked at a width of 5 mutations, while the involved decrease was at least 3-fold and reached up to approximately 100-fold. Thus, none of the trajectories to YQ:tggt was accessible by fixing mutations one-by-one in either of the two environments. While this analysis concerns only direct trajectories, i.e. without mutational reversions, allowing for reversions did not open up accessible trajectories. Overall, these data indicate that higher-order genetic interactions (i.e. epistasis involving multiple mutations) limit optimization of the *lac* regulatory phenotype in either of the two environments.

**Figure 2.**
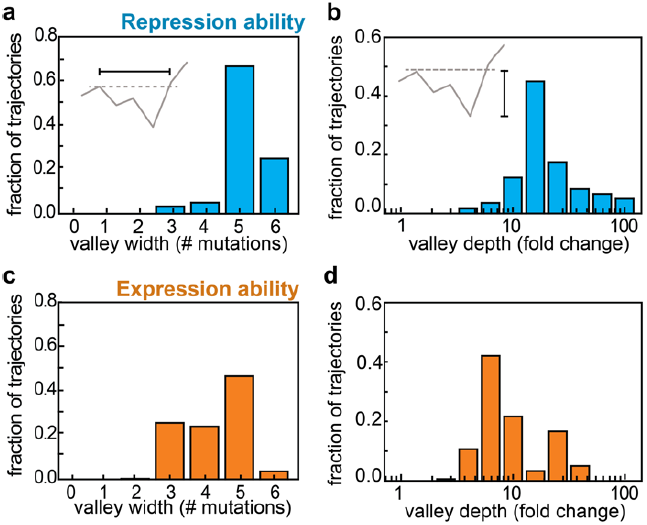
Genetic constraints in constant environments. (a) Valley width in repression ability (*R*), for all 720 mutational trajectories from MK:acca to YQ:tggt. Valley width equals the number of mutations required to regain a repression ability greater than or equal to the previous sub-optimum on a mutational trajectory. (b) Valley depth in repression ability. Valley depth is the largest decrease in repression ability along a mutational trajectory. (c) Valley width in expression ability (*E*). (d) Valley depth in expression ability. Welch’s t-tests were performed to determine the statistical significance of measured differences (Methods).

How does environmental variability affect these constraints? We first explored this question with individual trajectories starting with MK:acca. For instance, *R* could be increased through an operator mutation (MK:acca to MK:tcca) in the environment without ligand, but then remained trapped because the other mutations yielded no further improvements (Fig.3a and Supplementary Fig. 2). However, switching to the other environment opened up various trajectories that increased *E*, such as the repressor mutation MK:tcca to MQ:tcca. The system became trapped again on a suboptimum after a further increase in *E* (MQ:tcca to MQ:tgca). Switching back to the first environment now allowed escape, and provided access to YQ:tggt by increasing *R*. We found that a significant fraction of the direct trajectories (21%) became accessible in this manner (Supplementary Fig. 3). Thus, mutational pathways that failed to confer gradual optimization in either constant condition could do so when alternating between these same conditions.

**Figure 3.**
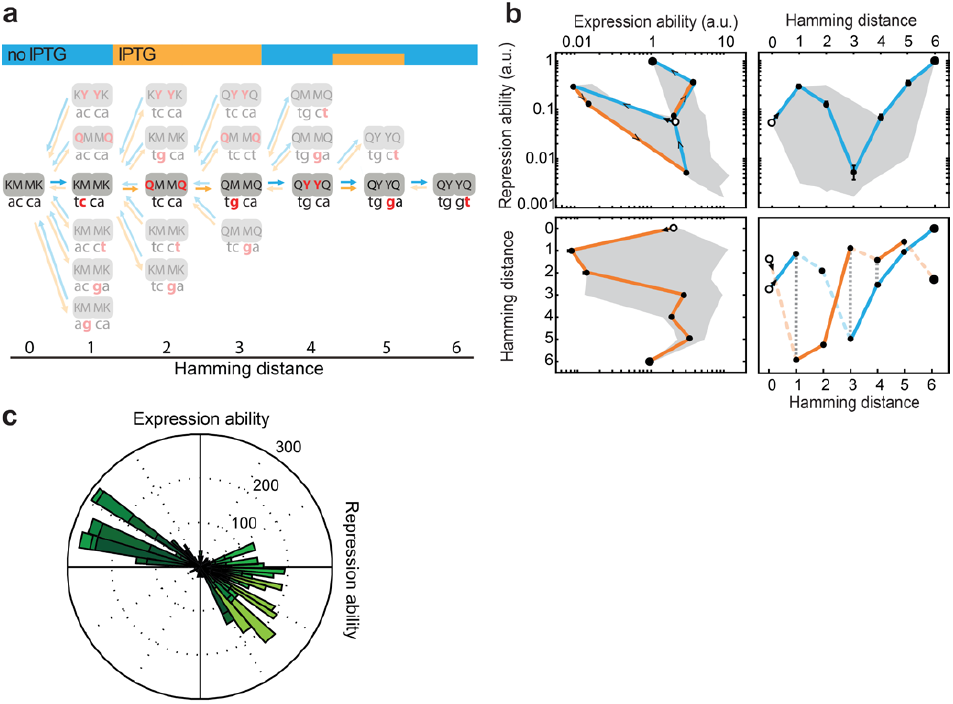
Escape from genetic constraint in fluctuating environments. (a) Mutational trajectory accessible by continuous improvements in a changing environment. Central genotypes correspond to the trajectory in Fig. 3b. Red indicates the mutated position. Forward arrows indicate mutations conferring increases in repression or expression ability, backward arrows indicate decreasing or neutral steps. Top color bar represents environmental changes that confer continuous improvements. Without these changes the system would be trapped at MK:tcca and MQ:tgca, where no further improvements are possible in the current environment. (b) Expression and repression ability along the trajectory indicated in panel (a). The trajectory starts at MK:acca (open circle) and ends at YQ:tggt (large filled circle). This trajectory contains a valley in repression ability (top right panel, blue line) and in expression ability (bottom left panel, orange line). Top left panel: Blue lines indicate mutations that confer improvements in repression ability, orange lines indicate mutations that confer improvements in expression ability. Bottom right panel: Dashed lines indicate mutations conferring deterioration. Grey dotted lines indicate environmental changes that allow escape from sub-optima. The grey area indicates the envelope of all trajectories. Error bars indicate standard errors. (c) Histogram of repression and expression ability effects of all mutations in all 720 trajectories. Most entries are in the top left and bottom right quadrants, indicating pervasive trade-off: mutations typically confer improvements in one and deterioration in the other environment. Mutations in the upper right quadrant are accessible in both environments. Color indicates Hamming distance, with lighter green corresponding to larger Hamming distance.

These findings cannot be understood from genotype x environment interactions that alter the magnitude of the mutational effect, as they would affect only the depth of constraining valleys. Rather, they can be explained by cross-environmental tradeoffs, in which increases in *R* occur at the expense of decreases in *E*, and vice versa, increases in *E* lead to decreases in *R* (Fig. 3b). Such tradeoffs were pervasive (85% of all mutations, Fig. 3c, Supplementary Fig. 4), and can be understood mechanistically. For example, a low but significant level of repression can be maintained in the presence of inducer through residual binding ^32^. We found that for several genotypes (22), the induced expression level was significantly lower than the highest measured level for the involved operator; consistent with the idea that induced repressors can reduce expression. Mutations that increase both this residual repression as well as the repression without inducer, as for instance achieved by overall increases in repressor-operator affinity, lead to opposite effects on *R* and *E*, and hence to cross-environmental trade-offs (Supplementary Fig. 4).

The cross-environmental trade-offs have important consequences for the relation between constraints in different conditions. We find multiple local optima for each of the two environments (3 in *R* and 13 in *E*), but none coincided at the same genotype. This feature allowed trajectories to repeatedly ‘surf’ ascending slopes in a ratchet-like mechanism: once trapped on a local optimum, waiting for an environmental change enabled repositioning on a new ascending slope. Thus, the fluctuating environment goes beyond providing a fluctuating selection strength, and rather opens up new trajectories by inverting the selective effect of mutations.

To assess the robustness of this evolutionary mode systematically for different conditions we extended a fixed-environment Markov approach ^33^ to include environmental fluctuations (Fig. 4a-c, Methods, Supplementary Text 1-6 and Supplementary Fig. 6 and 7). We consider a discrete-time Moran process in the strong selection weak mutation (SSWM) regime, in which trajectories can be of arbitrary length and allow mutational reversals (Methods) ^34, 35^. Consistent with the observed constraint in fixed conditions (Fig. 2), we found that the rate to access YQ:tggt from MK:acca (*k*_*e*_) was null for either constant environment (Fig. 4d). However, *k*_*e*_ was consistently above zero when the environmental fluctuation rate *k*_*f*_ was lower than the mutation rate *k*_*m*_ (Fig. 4d, blue and green lines) and maximized when *k*_*f*_ = *k*_*m*_, consistent with previous related work ^25^. This can be understood as follows: for *k*_*f*_ ≪ *k*_*m*_ the waiting time for an environment-triggered escape is long, while for *k*_*f*_ ≫ *k*_*m*_, there is an effective averaging over the two environments, which remains constraining.

**Figure 4.**
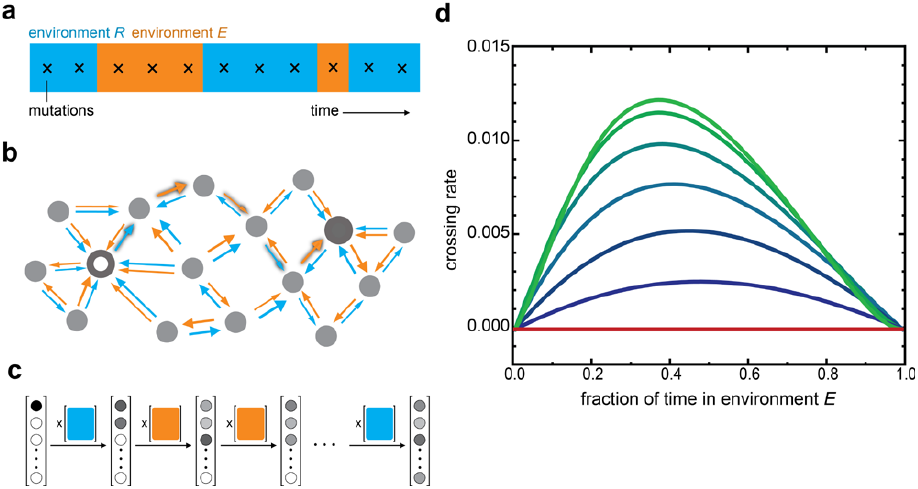
Landscape-crossing in stochastically alternating environments. (a) Environmental fluctuations and occurrence of mutations (crosses) (Supplementary Text 1). Environments *R* and *E* refer to the environment selecting for repression, resp. expression ability. (b) Schematic representation of genotype space. Large open and filled circles are start and end genotypes of mutational trajectories. Arrows indicate increasing repression ability (*R*, blue) or expression ability (*E*, orange); arrow thickness reflects magnitude and hence transition probability (Supplementary Text 2). Shadowed arrows indicate one possible path of continuous improvement from the initial to the final genotypes. The structure of the space is schematic and does not reflect the actual system. (c) Schematic depiction of the Markov chain method for computing crossing rates. The probability vector lists all *N* genotypes, with the grey-scale indicating the probability of populating a genotype at a given indicated time. Initially only the beginning genotype is populated. The *N* x *N* environment-dependent transition probability matrices (colour squares) reflect the arrows in panel B: a matrix entry at position i,j indicates the transition probability from genotype i to genotype j. Each matrix multiplication yields a novel probability of genotype-occupancy after a mutation occurred in a given environment. This illustration is schematic: we use (an infinite time limit) analytical solution for this process considering a range of possible scenario of environmental fluctuations. (d) Crossing rate as a function of fraction of time spent in each environment, for different environmental fluctuation rates. The unit of time is the time between two fixed mutations. Red line: environment dwell time << 1, meaning that the environment fluctuates much faster than the time between mutations. Top green line to bottom blue line: environment dwell time = 1, 2, 5, 10, 20, 50 (i.e. decreasing frequency of environmental fluctuation; Supplementary Text 3-6). The crossing rate is the inverse of the mean number of mutations that are necessary to cross the landscape. The maximum crossing rate is 0.17 (6^-1^, corresponding to six mutations).

Environmentally triggered escape was robust to changes in switching time between environments (Fig. 4d), as well as to the known non-linearity in the relation between *lac* expression and cellular growth rate (Supplementary Text 7, Supplementary Fig. 8) ^26, 36^. The mechanism broke down only when *lac* expression costs outweighed the benefits in the inducing environment, which is inconsistent with experimental observations ^26, 27^. Overall, these results showed that trade-offs can promote adaptive evolution in fluctuating conditions.

In summary, we report a mode of evolution in which phenotypes can break away from genetic constraints. In the presence of tradeoffs, environmental fluctuations can ‘ratchet’ phenotypes across valleys by continuous improvements. As environmental fluctuations and trade-offs are ubiquitous, this adaptive mechanism may well be relevant to a broad range of evolutionary transitions, and could have implications for clinical multi-drug protocols as well as evolutionary engineering.

## Acknowledgements

We thank D.M. Weinreich, J.A. de Visser, T. Paixão, J. Polechová, T. Friedlander and A.E. Mayo for reading and commenting on earlier versions of the manuscript, B. Houchmandzadeh, O. Rivoire and M. Hemery for discussions. Further, we kindly thank F.J.Poelwijk for sharing plasmid pCascade5 and pRD007 and Y. Yokobayashi for sharing plasmid pINV-110.

This work is part of the research program of the Foundation for Fundamental Research on Matter (FOM), which is part of the Netherlands Organization for Scientific Research (NWO).

### Statement of authorship

MGJdV and SJT, conception and design of experiments. MGJdV and VS performed experiments. MGJdV, AD and SJT analysed the data. AD formalised and analysed the mathematical model. MGJdV, AD and SJT wrote the manuscript.

